# Construction of habitat-specific training sets to achieve species-level assignment in 16S rRNA gene datasets

**DOI:** 10.1101/791574

**Authors:** Isabel F. Escapa, Yanmei Huang, Tsute Chen, Maoxuan Lin, Alexis Kokaras, Floyd E. Dewhirst, Katherine P. Lemon

**Affiliations:** Forsyth Institute (Microbiology), Cambridge, Massachusetts, USA; Department of Oral Medicine, Infection & Immunity, Harvard School of Dental Medicine, Boston, Massachusetts, USA; Alkek Center for Metagenomics & Microbiome Research, Department of Molecular Virology & Microbiology, Baylor College of Medicine, Houston, Texas, USA; Division of Infectious Diseases, Boston Children’s Hospital, Harvard Medical School, Boston, Massachusetts, USA; Section of Infectious Diseases, Department of Pediatrics, Texas Children’s Hospital and Baylor College of Medicine, Houston, Texas, USA

**Keywords:** training set, naïve Bayesian RDP Classifier, species-level taxonomy, 16S rRNA gene, V1-V3, habitat-specific database, microbiome, eHOMD, nasal, aerodigestive tract

## Abstract

**Background:** The low cost of 16S rRNA gene sequencing facilitates population-scale molecular epidemiological studies. Existing computational algorithms can parse 16S rRNA gene sequences to high-resolution Amplicon Sequence Variants (ASVs), which represent consistent labels comparable across studies. Assigning these ASVs to species-level taxonomy strengthens the ecological and/or clinical relevance of 16S rRNA gene-based microbiota studies and further facilitates data comparison across studies.

**Results:** To achieve this, we developed a broadly applicable method for constructing high-resolution training sets based on the phylogenic relationships among microbes found in a habitat of interested. When used with the naïve Bayesian Ribosomal Database Project (RDP) Classifier, this training set achieved species/supraspecies-level taxonomic assignment of 16S rRNA gene-derived ASVs. The key steps for generating such a training set are: 1) constructing an accurate and comprehensive phylogenetic-based, habitat-specific database; 2) compiling multiple 16S rRNA gene sequences to represent the natural sequence variability of each taxon in the database; 3) trimming the training set to match the sequenced regions, if necessary; and 4) placing species sharing closely related sequences into a supraspecies taxonomic level to preserve subgenus-level resolution. As proof of principle, we developed a V1-V3 region training set for the bacterial microbiota of the human aerodigestive tract using the full-length 16S rRNA gene reference sequences compiled in our expanded Human Oral Microbiome Database (eHOMD). We also overcame technical limitations to successfully use Illumina sequences for the 16S rRNA gene V1-V3 region, the most informative segment for classifying bacteria native to the human aerodigestive tract. Finally, we generated a full-length eHOMD 16S rRNA gene training set, which we used in conjunction with an independent PacBio Single Molecule, Real-Time (SMRT)-sequenced sinonasal dataset to validate the representation of species in our training set. This also established the effectiveness of a full-length training set for assigning taxonomy of long-read 16S rRNA gene datasets.

**Conclusion:** Here, we present a systematic approach for constructing a phylogeny-based, high-resolution, habitat-specific training set that permits species/supraspecies-level taxonomic assignment to short- and long-read 16S rRNA gene-derived ASVs. This advancement enhances the ecological and/or clinical relevance of 16S rRNA gene-based microbiota studies.

## BACKGROUND

In microbiota studies of most ecosystems and/or habitats, achieving species- or strain-level identification of constituents improves the ecological and/or clinical relevance of the results compared with genus-level identification. For example, species-level identification is often critically important for host-associated microbial communities because these communities frequently include commensal and pathogenic species of the same genus, e.g., [1, 2], with the caveat that strain-level variation in pathogenicity also exists, e.g., [3]. Additionally, some microbial genera include species that are site specialists and inhabit distinct niches of a given environment [4]. High-throughput close-to-full-length 16S rRNA gene sequencing (e.g., circular consensus sequences from PacBio Single Molecule, Real-Time (SMRT) sequencing) and metagenomic <u>w</u>hole <u>g</u>enome <u>s</u>equencing (WGS) hold promise for species- and strain-level microbiota studies. However, the easier accessibility, and lower cost, of 16S rRNA gene short-read sequencing makes population-scale (i.e., thousands of samples) molecular epidemiological studies of bacterial microbiota of humans, other animals, plants, and the environment broadly feasible now. A caveat to this is that the majority of published short-read 16S rRNA gene sequencing studies use read clustering at a percent similarity that constrains resolution to the genus level, i.e., 97% identity. Indeed, recent reviews on best practices and benchmarking for 16S rRNA gene microbiota studies focus on genus-level operational taxonomic unit (OTU) analysis, e.g., [5-7]. However, newer algorithms, many of which are not based on similarity thresholds, allow single-nucleotide resolution and can parse 16S rRNA gene short-read sequences into species- or strain-level resolution phylotypes, usually called <u>a</u>mplicon <u>s</u>equence <u>v</u>ariants (ASVs) (e.g., MED (Minimal Entropy Decomposition) [8, 9], DADA2 (Divisive Amplicon Denoising Algorithm) [10, 11] and UNOISE [12, 13], among others [14, 15]). Another limitation of short-read 16S rRNA microbiome studies is that the choice of the 16S rRNA gene region(s) sequenced places an upper bound on the degree of species-level resolution that is achievable within a dataset [16-19]. Therefore, it is critical to determine which regions provide the most information for distinguishing bacterial species that are common to that specific ecosystem. For the habitats within the human aerodigestive tract, i.e., the nasal passages, sinuses, throat, mouth, esophagus, as well as the lower respiratory tract, we previously showed that many more taxa are distinguishable at the species level with the V1-V3 region than with the commonly used V3-V4 region of the 16S rRNA gene [20]. Therefore, we have developed a method that, by combining the “reusability, reproducibility and comprehensiveness” of ASVs [11, 13] and the selection of highly informative regions of the 16S rRNA gene, maximizes 16S rRNA gene short-read sequencing potential to achieve sub-genus level resolution taxonomic assignment.

Microbial databases encompassing broad phylogenetic diversity, such as SILVA [21, 22], RDP [23] and Greengenes [24], serve the key role of being applicable to myriad different habitats. However, this valuable breadth comes with the trade-off of inclusion of taxonomically misannotated 16S rRNA gene sequences. For example, Edgar estimated annotation error rates as high as ∼10-17% in these comprehensive databases [25]. SILVA and RDP continue to undergo regular updates and contain a broadly expansive and comprehensive record of 16S rRNA gene sequences from all explored habitats, whereas Greengenes was last updated in 2013. For habitats that have yet to be deeply interrogated, the access to this breadth outweighs the risk of misclassification due to annotation error. However, once a habitat is sufficiently explored through sequencing, creation of a habitat-specific database enables accurate fine-level phylogenetic resolution for taxonomic assignment to ASVs [20, 26-35]. Existing habitat-specific databases are constructed with different methods and can be used to assign taxonomy via different approaches, as in the following three examples. First, there are stand-alone habitat-specific databases consisting of curated collections of close-to-full-length 16S rRNA gene sequences compiled both from other repositories and by generating new sequences from the habitat of interest, e.g., eHOMD for the human aerodigestive tract [20, 26, 36], HITdb for the human gut [29] and RIM-DP for rumen [28]. Second, custom addition of compiled sequences from a specific habitat of interest can be used to augment a broad general database, e.g., HBDB for honey bee [27], DictDB for termite and cockroach gut [33], SILVA19Rum for rumen [35] and MiDAS for activated sludge [30, 32]. Third, both a general and a habitat-specific database can be combined in the same pipeline, e.g., for human-associated genera with pathogenic members [37], for freshwater (FreshTrain with the TaxAss workflow [34]) and the multistage blastn workflow with eHOMD [38]. Many of these databases are formatted for use as training sets to train classifier algorithms for taxonomy assignment.

The naïve Bayesian RDP Classifier [39] is one of several effective algorithms for assigning taxonomy. (For benchmarking and comparison with other methods, see [40-43]). This type of supervised learning algorithm requires a training set, which is a set of input-output examples to learn a function that can be used to make predictions [44]. In this case, sequences are input and taxonomic assignments are output. Properly formatted versions of the broad 16S rRNA gene databases SILVA, RDP and Greengenes are available to train the most popular implementations of the naïve Bayesian RDP Classifier. The quality of the training set strongly influences taxonomic assignment and habitat-specific training sets have been developed to increase accuracy of taxonomic assignments [27, 33, 40-42, 45]. However, the resolution of available training sets is mostly limited to the genus level. An exception is the manually curated subset of the Greengenes database corresponding to 89 clinically relevant bacterial genera that was used to assign species-level taxonomy of full-length 16S rRNA gene sequences of clinical isolates [46]. Notwithstanding, species-level taxonomy assignment of short-read 16S rRNA gene datasets remains a challenge.

We hypothesized that we could develop a method to rapidly generate a habitat-specific training set to leverage the strength of the naïve Bayesian RDP Classifier to consistently achieve species- or supraspecies (i.e., subgenus)-level taxonomic assignment of ASVs starting from a locally comprehensive, phylogeny-based, high-resolution (i.e., separated at ≥ 98.5% identity) set of curated reference sequences with distinct taxonomic names. Here, we show that the use of the naïve Bayesian RDP Classifier with a training set in which each taxon is represented by a collection of highly similar sequences that captures the natural variability of each species resulted in accurate species-level taxonomic assignment of short-read and long-read sequences of the 16S rRNA gene. This represents a methodological advancement. Our systematic approach for generating training sets is applicable to any ecosystem/habitat of interest and is summarized in **Figure 1**. This approach requires compiling high-quality close-to-full-length 16S rRNA gene datasets from a habitat (**Figure 1A**). These compiled datasets are then used to identify curated reference sequences to build a 16S rRNA gene database (**Figure 1B**) from which training sets for that habitat are derived (**Figure 1C**).

**Figure 1.**
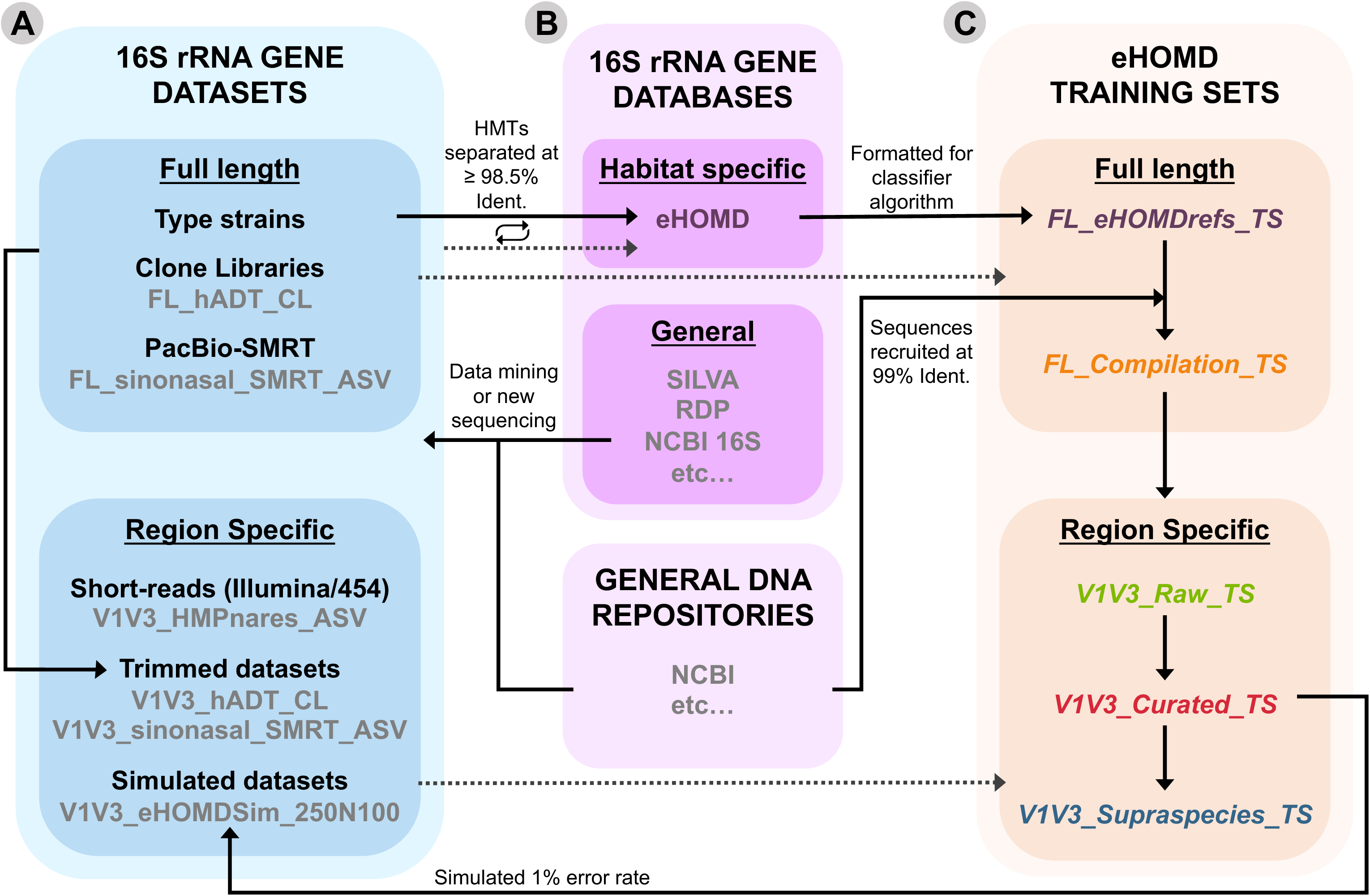
Relationships between the datasets, databases and training sets in constructing training sets for a specific habitat: the human aerodigestive tract. **A)** Datasets gathered from public repositories or obtained by sequencing of new samples are used to explore the 16S rRNA gene diversity of the habitat of interest. These include both 16S rRNA full-length sequences and region-specific short-read sequences used for method validation or benchmarking. **B)** A curated habitat-specific full-length 16S rRNA gene reference database is assembled and expanded in an iterative way by selecting from those datasets in A representative sequences for both named and as-yet unnamed or uncultivated species (i.e., HMTs in eHOMD), and placing them in a phylogenetic tree (See Figure 1 in [20]). **C)** Training sets are derived from the taxonomical hierarchy of the habitat-specific database and enhanced by the following steps: compiling multiple 16S rRNA gene sequences to represent the natural sequence variability of each taxon, trimming the training set to match the sequenced region/s, and placing species sharing closely related sequences into a supraspecies taxonomic level. To enhance clarity, the *names of training sets* appear in bold italics whereas the **names of datasets** appear in bold throughout this paper. Datasets in grey are the specific examples used for the construction of the eHOMD derived training sets described here. Solid arrows indicate where the sequences described come from and dotted arrows indicate when datasets were used for validation or benchmarking.

To test our hypothesis, we developed and validated short- and long-read training sets for the microbiota of the human aerodigestive tract (mouth, nasal passages, sinuses, throat, and esophagus) using our expanded Human Oral Microbiome Database (<u>eHOMD</u>). This database was originally created and later expanded to serve as a resource for the community of investigators generating datasets to study habitats within the human aerodigestive tract [20, 26, 36]. In addition to 16S rRNA gene reference sequences (eHOMDrefs), it also includes genomic and proteomic data. (It also works well for the lower respiratory tract [20].) The lack of proper taxonomical representation in traditional databases is a challenge in predicting taxonomic assignments [43]. A strength of the eHOMD is that, by placing 16S rRNA gene reference sequences for each Human Microbial Taxon (HMT) on a phylogenetic tree (http://ehomd.org/index.php?name=HOMD&show_tree=_), as-yet unnamed or uncultivated species are defined based on sequence identity and added to the phylogeny using a provisional naming scheme that permits taxonomic assignment for cross-study comparison [26]. Also, sequences that are misnamed in other databases are easily identified and given a correct designation in eHOMD. Another key strength of eHOMD is that it is locally comprehensive often allowing approximately 95% of sequences from V1-V3 aerodigestive tract datasets to be assigned to the taxonomy [20].

## RESULTS

### Compiling closely-related sequences for each taxon in a training set improves the accuracy of species-level taxonomic classification

Genus-level taxonomic assignment is not an inherent limitation of the naïve Bayesian RDP Classifier. Rather, taxonomic assignment to 16S rRNA gene short reads is limited by both the resolution to which sequences in datasets are parsed and by the nature of the training set used. The former is overcome by using approaches such as oligotyping/MED [8, 9], DADA2 [10, 11] or unoise2 [12, 25] to parse sequence variants at high resolution. We hypothesized that the limitations inherent in training sets could also be overcome.

The naïve Bayesian RDP Classifier algorithm indicates that a training set with a larger number of sequences representing each taxon will result in more confident taxonomic assignment [39]. Based on the conditional probability for a member of a taxon (T), the higher the occurrence of a given distinguishing “k-mer” (word or *w_i_*) in the training set, the greater the confidence with which assignment of that taxon is made, i.e., more sequences can be classified unambiguously. Thus, as the number of sequences (M) for each taxon in the training set increases (**Figure 2A** **vs. 2B**), the number of accurate assignments should increase. (See **Additional File 1** for a more detailed explanation.) In support of this, Werner and colleagues found more sequence counts result in more assignments [45]. Therefore, to systematically increase M, we used each of the reference sequences in eHOMD, i.e., the eHOMDrefs, as bait to capture closely matching, publicly available sequences and combined the resulting compilation of sequences for each taxon into a close-to-full-length compilation training set (***FL_Compilation_TS***) that reflects the currently known 16S rRNA gene sequence variability for each taxon, both natural and sequencing-error derived (**Additional File 2**).

**Figure 2.**
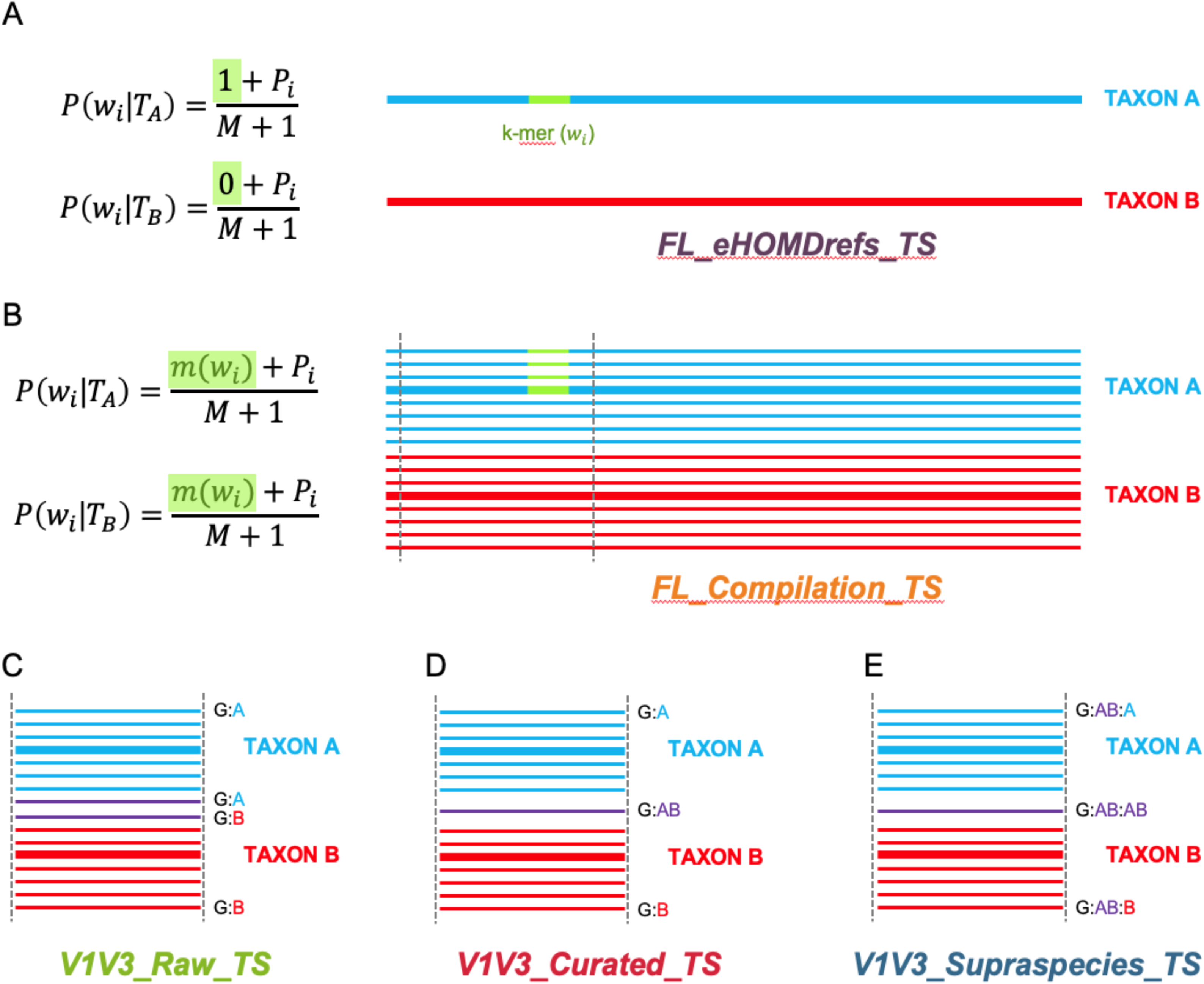
Schematic representation of the steps to generate sequential habitat-specific training sets. (**a**) The ***FL_eHOMDrefs_TS*** training set contains all full-length eHOMDrefs (thick lines) from eHOMDv15.1 together with their respective taxonomic assignment. When only one read represents each taxon (M=1) a given distinguishing k-mer (green fragment) can only be either present (1) or absent (0). (**b**) A higher number of sequences per taxon (M) allows for better resolution on the assignment, with the presence of a given distinguishing k-mer (green fragment) across each cluster of reads (w_*i*_) being represented as a proportion out of the total number of reads in that taxon (M). Therefore, to better represent the known sequence diversity of the 16S rRNA gene(s) for each taxon, the training set ***FL_Compilation_TS*** includes clusters of sequences (thin lines) recovered from the NCBI nonredundant nucleotide (nr/nt) database that matched with 99% identity and ≥ 98% coverage (see methods) to each eHOMDref (thick line). (**c**) The training set ***V1V3_Raw_TS*** is a V1-V3 trimmed version of the ***FL_Compilation_TS*** training set. The schematic illustrates how trimming to this region leads to identical reads (purple lines) having two different taxonomic designations. Here, G is genus and species are labeled as A or B. (**d**) To construct the ***V1V3_Curated_TS*** training set, identical V1-V3 sequences in the ***V1V3_Raw_TS*** training set were collapsed into one. If identical sequences came from more than one taxon (purple), species-level names of all taxa involved were concatenated (AB). (**e**) The ***V1V3_Supraspecies_TS*** training set includes the same sequences that the ***V1V3_Raw_TS*** training set; however, the headers in the fasta file include the supraspecies taxon (AB) as an extra level between the genus (G) and species taxonomic levels (A, B or AB), as illustrated here.

To evaluate the performance of the ***FL_Compilation_TS*** and for further method optimization, we created the simulated dataset **V1V3_eHOMDSim_250N100** by introducing a 1% error rate in a V1-V3 trimmed version of our training set sequences. (See Methods for details and **Additional File 3** for the dataset itself.) This simulated dataset aims to mimic real sequencing short-reads similar to the ones that can be obtained with our method for achieving highly informative 16S rRNA gene V1-V3 region sequencing data using Illumina MiSeq, which is detailed in **Additional File 4**. We then assessed the percentage of reads in the simulated dataset **V1V3_eHOMDSim_250N100** that classified at the species-level at incremental bootstrap values from 50 to 100 using the training set ***FL_Compilation_TS*** (**Figure 3A**, orange bars) compared with the training set ***FL_eHOMDrefs_TS*** (**Figure 3A**, purple bars), which consists of only the eHOMDrefs. Doing this, we observed an increase in the percentage of reads that classified at species-level with the compilation TS except at a bootstrap value of 100. We postulate that the additional sequences classified with the ***FL_eHOMDrefs_TS*** TS at a bootstrap of 100 are misclassified sequences. A higher rate of misclassification is expected with the training set ***FL_eHOMDrefs_TS***, since it includes only a few representative sequences for each taxon. Therefore, we next assayed for any misclassifications. At each bootstrap threshold, the percentage of reads that were misclassified (i.e., reads for which the assigned taxonomic identity was different than the known identity of the original sequence from which the simulated read was derived) was at least 50% lower using the training set ***FL_Compilation_TS*** (**Figure 3B**, orange line) than with the training set ***FL_eHOMDrefs_TS*** (**Figure 3B**, purple line). Thus, classification of the dataset **V1V3_eHOMDSim_250N100** showed a reduced error rate and increased confidence level when using a training set consisting of a compilation of closely related sequences rather than one consisting of only one or a few reference sequences for each taxon.

**Figure 3.**
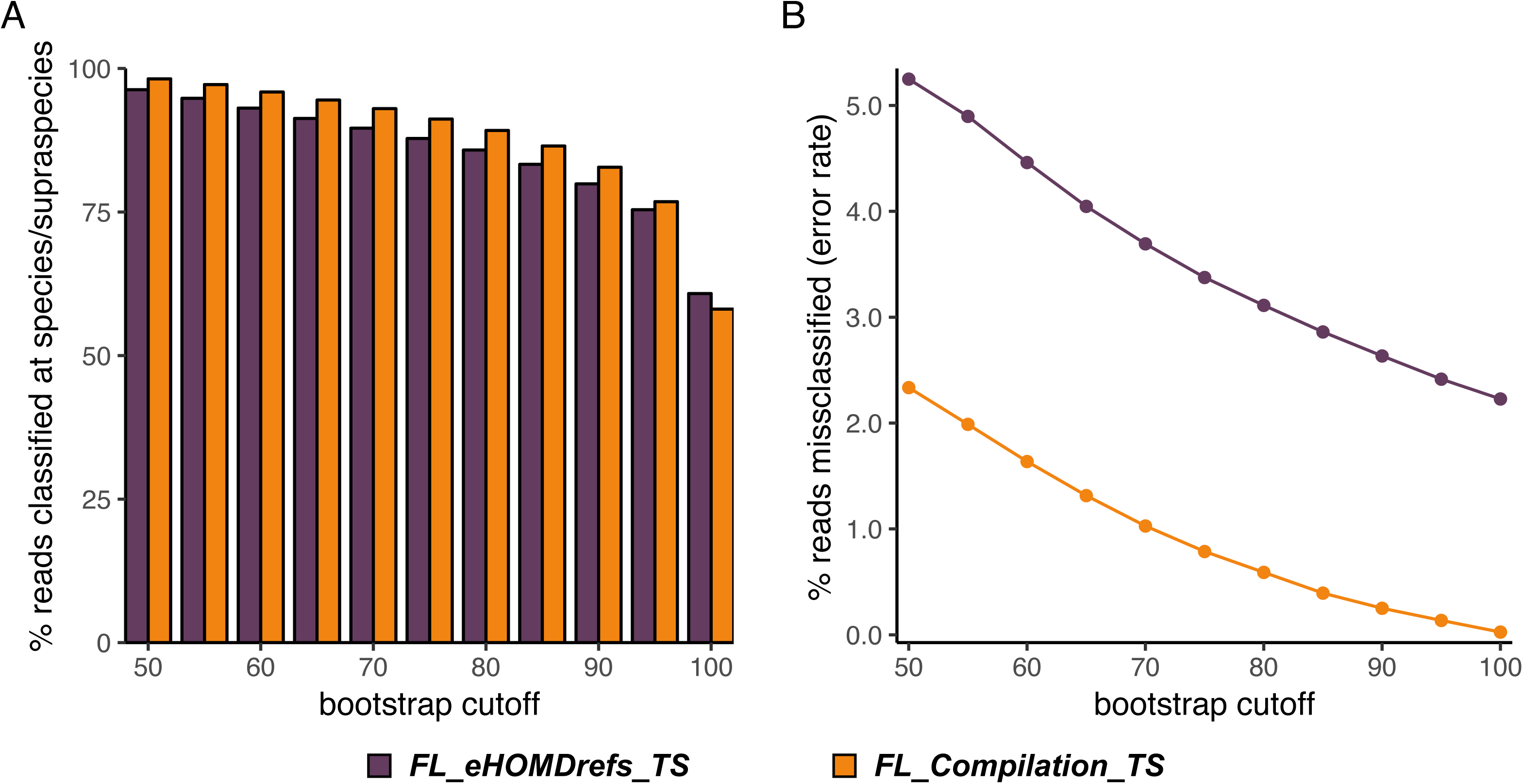
The *FL_Compilation_TS* training set provides higher classification percentages with a lower error rate. The naïve Bayesian RDP Classifier was used with bootstrap values ranging from 50 to 100. (**a**) The percentage of eHOMD-derived simulated reads classified using the ***FL_eHOMDrefs_TS*** training set (purple) versus the ***FL_Compilation_TS*** training set (orange). (**b**) The percentage of classified reads that were misclassified (i.e., reads for which the assigned taxonomic identity was different than the known identity of the original sequence from which the simulated read was derived).

### Moving towards an appropriate short-read fragment training set

The training set ***FL_Compilation_TS*** consisted of close-to-full-length 16S rRNA gene sequences. We hypothesized that the presence of k-mers outside the V1-V3 region in the training set might lead to misclassifications when using it with a V1-V3 region dataset. Supporting this, when using a training set based on a large general database, trimming reference sequences to match the sequenced region increases the number of sequences that are assigned taxonomy [45]. Therefore, we trimmed the sequences in the training set ***FL_Compilation_TS*** to cover only the V1-V3 region generating training set ***V1V3_Raw_TS*** (**Figure 2C**). Using this there was no gain in the percentage of reads in the simulated dataset **V1V3_eHOMDSim_250N100** classified to the species-level (**Figure 4A**, green bar). Moreover, we observed an increase in the percentage of misclassified reads (**Figure 4B**, green line), i.e. the assignment accuracy decreased. Therefore, we next determined why the use of appropriate short-read fragments in this training set paradoxically decreased confidence and accuracy of species-level assignment and resolved the issue.

**Figure 4.**
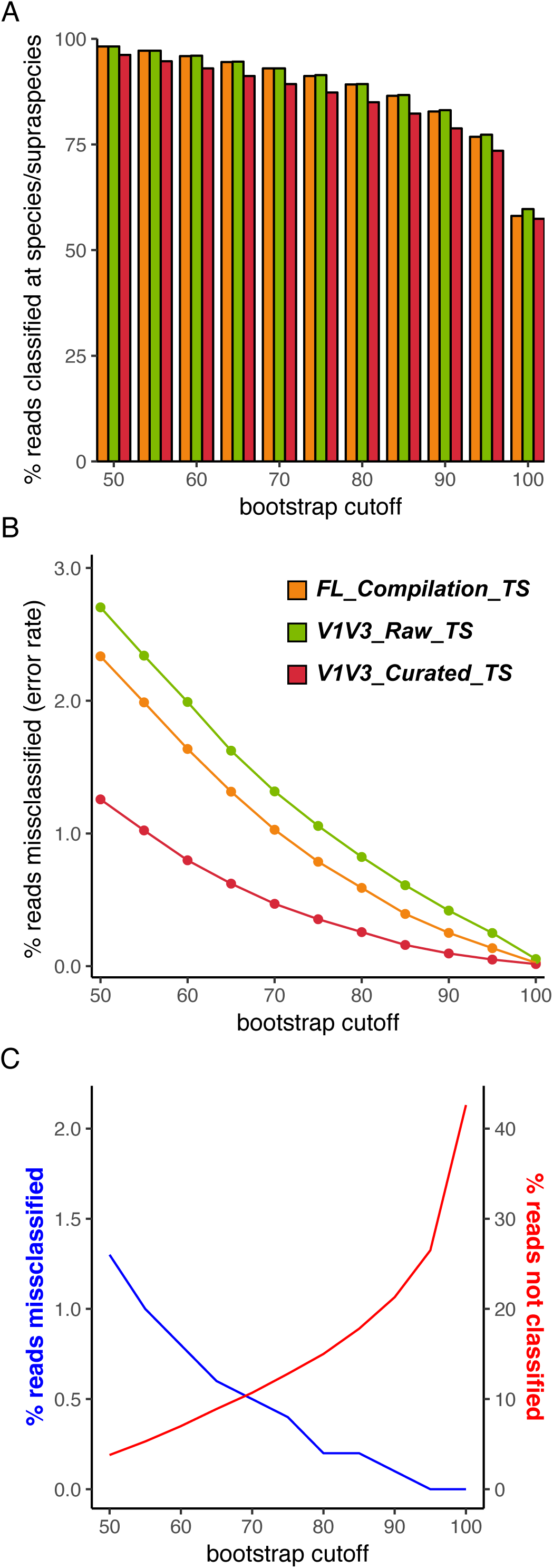
Trimming the training set to the specific sequenced region further reduces the error rate. (**a**) The percentage of eHOMD-derived simulated reads classified at species level using the ***FL_Compilation_TS*** (orange) training set compared to subsequent trimmed versions ***V1V3_Raw_TS*** (green) and ***V1V3_Curated_TS*** (red). (**b**) The percentage of classified reads that were misclassified with each of these three training sets. The naïve Bayesian RDP Classifier was used with bootstrap values ranging from 50 to 100. (**c**) This graph, which is specific to the eHOMD training set construction (**V1V3_eHOMDSim_250N100** dataset), indicates how researchers can determine the bootstrap value to use with the naïve Bayesian RDP Classifier by deciding an acceptable level of the % of reads misclassified (blue line; e.g., 0.5%) and/or of the % of reads that are not classified (red line).

### Combining closely related, indistinguishable taxa into supraspecies decreases the error rate for a short-read training set

Considering the possible explanations for the above paradox, we realized that taxa with distinct full-length 16S rRNA gene sequences can have identical V1-V3 sequences. In silico, using only V1-V3, 37 of the ∼770 species-level taxa in eHOMD are no longer distinguishable from at least one other species at 100% identity [20]. Therefore, we hypothesized that the observed loss in accuracy using training set ***V1V3 _Raw_TS*** was due to identical sequences with more than one species name, e.g., *Veillonella parvula* and *Veillonella dispar* [20]. To solve this problem, we removed duplicate sequences and assigned a combined name, i.e., a supraspecies name such as *Veillonella* parvula_dispar, to the remaining unique sequence (**Figure 2D**). This resulted in the training set ***V1V3_Curated_TS***, which showed improved accuracy compared with both the ***FL_Compilation_TS*** and the ***V1V3_Raw_TS*** (**Figure 4B**, red line). However, this improvement came at a cost of a 0.7 to 4.4 % decrease in the reads assigned supraspecies- or species-level taxonomy at each bootstrap threshold (**Figure 4A**, red bar). This trade-off can be illustrated by graphing at each bootstrap threshold the percentage of reads from the simulated **V1V3_eHOMDSim_250N100** dataset that were misclassified versus confidently assigned species-level taxonomy using the ***V1V3_Curated_TS*** with the naïve Bayesian RDP Classifier (**Figure 4C**).

### Maximizing performance of the naïve Bayesian RDP Classifier for subgenus-level taxonomic assignment of short-read sequences requires both insertion of supraspecies as a taxonomic level and setting a threshold bootstrap

Two steps were required to reap the benefits of using the supraspecies definition. First, formally inserting supraspecies as a taxonomic level between genus and species in the name header for each sequence in the training set, which yielded the training set ***V1V3_Supraspecies_TS*** (**Figure 2E; Additional File 5**). Second, establishing the bootstrap cut-off at which a sequence is not assigned at the species level so that the naïve Bayesian RDP Classifier will then default to the supraspecies level, rather than defaulting to genus, allowing for a higher percentage of reads to be assigned (**Figure 5A**, blue bar). This latter step preserves subgenus level information encoded in the ASVs. This choice inevitably involves a trade-off between accuracy and percentage of reads classified below genus level, e.g., see **Figure 4C**. For our purposes, we chose a conservative bootstrap value of 70 (**Figure 5B**, blue line). With the simulated data, for which the truth is known, this gave an error rate of around 0.05%. Of note, although one common bootstrap setting for the naïve Bayesian RDP Classifier is 50, we use of a more conservative value for species-level taxonomic assignment with the ***V1V3_Supraspecies_TS***.

**Figure 5.**
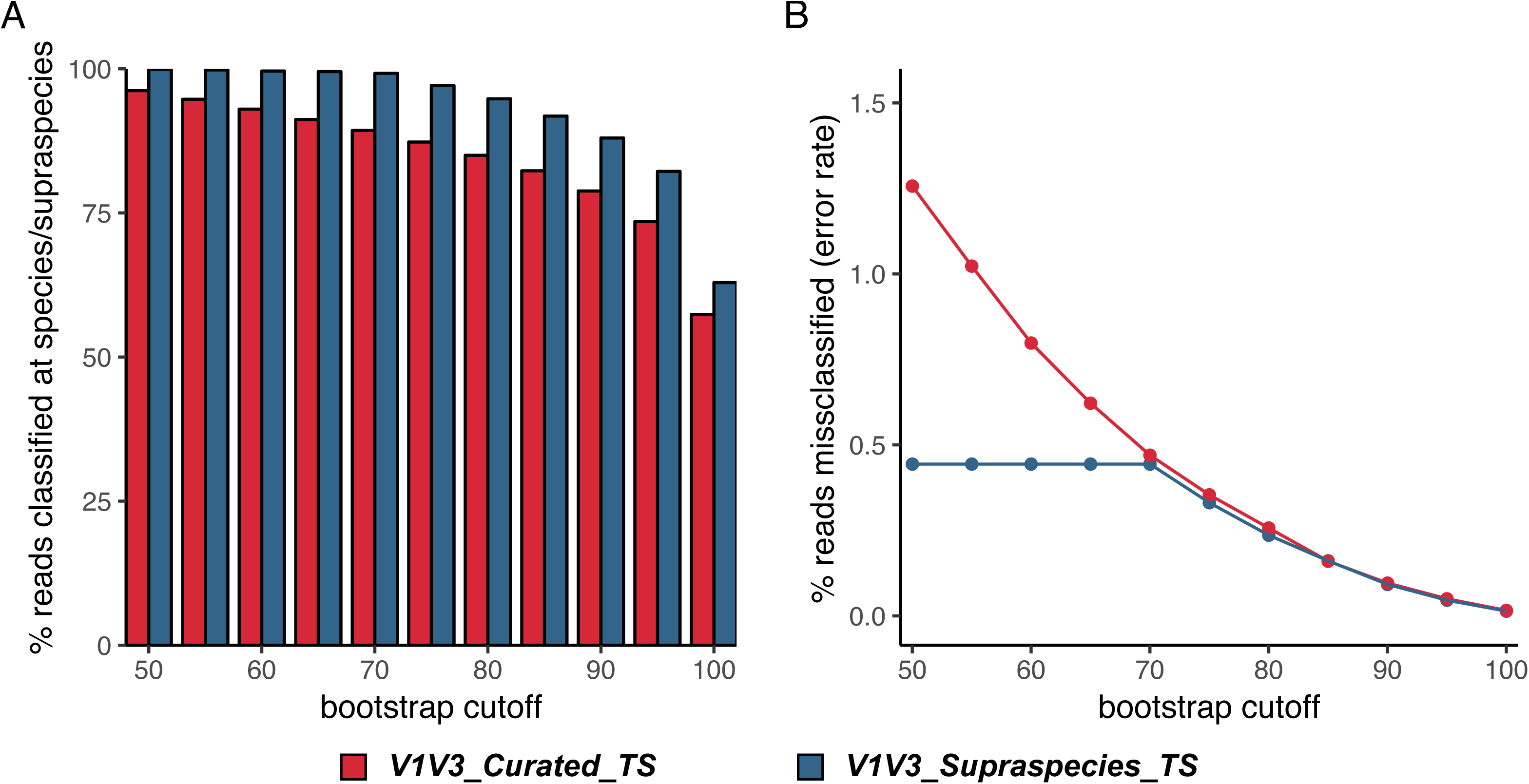
Addition of a supraspecies level to the training set increases the percentage of classified reads. (**a**) The percentage of eHOMD-derived simulated reads classified at species/supraspecies level using the ***V1V3_Curated_TS*** training set (red) versus the ***FL_Supraspecies_TS*** training set (blue). (**b**) Percentage of classified reads that were misclassified with each of these TS. The naïve Bayesian RDP Classifier was used with bootstrap values ranging from 50 to 100.

### Using an independent sinonasal dataset validated the performance of the eHOMD training set

We next validated the performance of the eHOMD training set against a dataset of close-to-full-length 16S rRNA gene sequences from sinonasal samples. Very importantly, these sequences are independent of the sequences recruited to build the training set. Earl and colleagues performed PacBio SMRT sequencing of close-to-full-length 16S rRNA genes from multiple sinonasal samples from each of 12 adults without sinus inflammation who were undergoing removal of a pituitary adenoma [47]. Because this dataset was deposited in the SRA, these sequences were not present in the NCBI nr/nt sequence repository (from which sequences were recruited for the eHOMD training set) and therefore not used in the generation of the eHOMD training set. We used the DADA2 pipeline for PacBio SMRT reads [48] to denoise these sequences and identify the relative abundance of each ASV in this dataset (**FL_sinonasal_SMRT_ASV; Additional File 6**). Next, we followed a three-pronged approach to assess the validity of the eHOMD training set for use with this fully independent sinonasal dataset.

First, we compared the best match by blastn, as a proxy for taxonomic assignment for these ASVs, using eHOMD compared to using NCBI 16S Microbial to identify any taxa in the dataset that might be missing in eHOMD, and thus the training set (**Table 1**; **Additional File 7A**). A detailed comparison (**Additional File 7B**), showed that 50 ASVs, corresponding to 11.1% of the total reads, were differentially assigned by blastn (98.5% identity cutoff) against eHOMDv15.1 versus against NCBI 16S Microbial. Of these 50 ASVs, 33 were assigned to taxa with an HMT (Human Microbial Taxon) # as a provisional species designation. The majority (17/33) were assigned to *Peptoniphilus* sp. HMT-187, which corresponded in the NCBI 16S Microbial blastn output to *Peptoniphilus lacydonensis* and accounted for 7.6 % of the total reads. Similarly, *Peptoniphilus* sp. HMT-187 was 11th in total reads across the nostril samples of 210 HMP (Human Microbiome Project) participants [20]. The new species name *Peptoniphilus lacydonensis* was published in 2018 and is formally recognized [49]; therefore, it is now linked to HMT-187 in <u>eHOMD</u>. Of the remaining 17 ASVs, 8 were mismatches between closely related *Streptococcus* species. Only 6 ASVs were nonassigned (NA) using eHOMD, thus, eHOMD included the vast majority (99.7%) of taxa present in the Earl-Mell sinonasal dataset. We note that based on recent additions both eHOMD and NCBI 16S Microbial allowed correct assignment of ASVs to *Lawsonella clevelandensis*, which previously might be misidentified by blastn best match as a Dietzia species, and to *Neisseriaceae* G-1 bacterium HMT-327, which before receiving a stable taxonomic designation in eHOMD might be misidentified by blastn best match as a *Snodgrassella* species [20].

**Table 1.** The eHOMD training sets performed well for species/supraspecies taxonomy assignment of an independent long-read 16S rRNA gene sinonasal dataset.

|  | # ASVs | # Reads | % ASVs | % Reads |
| --- | --- | --- | --- | --- |
| NCBI 16S Microbial blastn | 178 | 140,455 | 87.3 | 97.3 |
| eHOMD blastn | 188 | 143,274 | 92.2 | 99.3 |
| eHOMD *FL_Compilation_TS* | 194 | 142,843 | 95.1 | 99.0 |
| eHOMD V1V3_Supraspecies_TS | 201 | 144,279 | 98.5 | 99.9 |

Second, we compared species-level assignment of the dataset **FL_sinonasal_SMRT_ASV** using blastn against eHOMD versus the naïve Bayesian RDP Classifier with the full-length eHOMD training set ***FL_Compilation_TS***. Less than 2% (1.91%) of the total reads in the dataset **FL_sinonasal_SMRT_ASV**, which are represented by 25 ASVs, were differentially assigned species-level taxonomy by these approaches (**Table 1**; **Additional File 7A**). There was no clear pattern to this differential assignment (**Additional File 7C**). This served to validate the use of the full-length eHOMD training set ***FL_Compilation_TS*** and demonstrated that our approach is an effective method for assigning species-level taxonomy to close-to-full-length 16S rRNA gene sequences such as those derived from PacBio SMRT sequencing.

Third, based on the above, we used naïve Bayesian RDP Classifier-derived assignments with the training set ***FL_Compilation_TS*** as a proxy for the true species representation in the dataset **FL_sinonasal_SMRT_ASV**. We then used these to assess the performance of the training set ***V1V3_Supraspecies_TS*** for assigning species-level taxonomy to the V1-V3 region sequences of the dataset **FL_sinonasal_SMRT_ASV** (**V1V3_sinonasal_SMRT_ASV; Additional File 8**). There were seventeen V1-V3 ASVs that were differentially assigned, corresponding to 1.64% of the reads (**Table 1**; **Additional File 7D**). Of these 17 differentially-assigned ASVs, 5 belonged to a supraspecies, a level not represented in the full-length training set ***FL_Compilation_TS***. Another 4 ASVs, all of which were present at very low relative abundance, had full-length ASVs assigned to *Anaerococcus tetradius*, whereas the V1-V3 region was assigned to *Anaerococcus octavius*, all with bootstrap values ≤ 92 (**Additional File 7A**). In contrast, when 16S rRNA gene sequences were parsed into ASVs based on their full-length and then cut to V1-V3, there was little to no species-level ambiguity (**Additional File 7D**). Thus, the ambiguity observed with V1-V3 16S rRNA short reads most likely occurs at the level of parsing into ASVs, due to fewer informative sequences, rather than at the assignment step with the naïve Bayesian RDP Classifier and the eHOMD training set.

### The eHOMD training set outperforms both the SILVA and RDP training sets

Having validated the use of the eHOMD training set with an independent 16S rRNA gene sinonasal dataset (**Table 1**), we next compared the use of our taxonomic assignment approach with other currently available pipelines that are coupled with the RDP or SILVA databases. We used three different datasets for this: 1) we generated a V1-V3 dataset derived from a collection of <u>h</u>uman <u>a</u>ero<u>d</u>igestive tract 16S rRNA gene clone libraries (**V1V3_hADT_CL; Additional File 9**); 2) the HMP 16S rRNA gene V1-V3 454-sequenced nostril dataset that we previously analyzed [20]; and 3) the close-to-full-length ASVs of the **FL_sinonasal_SMRT_ASV** dataset. We then assigned genus-level taxonomy to all of these using the naïve Bayesian RDP Classifier with a bootstrap threshold of 70 coupled with three different training sets: an eHOMD training set (V1-V3 or FL), RDP16 or SILVA132 (the latter two from https://benjjneb.github.io/dada2/training.html). The eHOMD training set resulted in a larger percentage of reads assigned to a specific genus for each dataset; however, all three training sets resulted in genus-level assignment of >90% of the sequences (**Table 2**). In contrast, a striking difference emerged between the different workflows when assigning taxonomy to species level. Our method of coupling the naïve Bayesian RDP Classifier with an eHOMD training set showed superior performance in terms of the percentage of reads classified compared to the exact string match method that is currently implemented in the DADA2 R Package in conjunction with SILVA132 or RDP16 (**Table 2**). As would be expected, the exact match algorithm assigned a higher percentage of close-to-full-length ASVs than V1-V3 region ASVs to taxonomy. The caveat being that these comprehensive databases have an estimated annotation error rates as high as ∼10-17% [25]. For the V1-V3 region ASVs, it also performed much better with the V1-V3 sequences in the V1-V3 human aerodigestive tract clone library dataset than with the HMP V1-V3 dataset. We speculate this occurred because the close-to-full-length sequences from the human aerodigestive tract clone library (**V1V3_hADT_CL**) dataset are part of both the RDP and SILVA databases, whereas the HMP V1-V3 454 sequences are not.

**Table 2.**
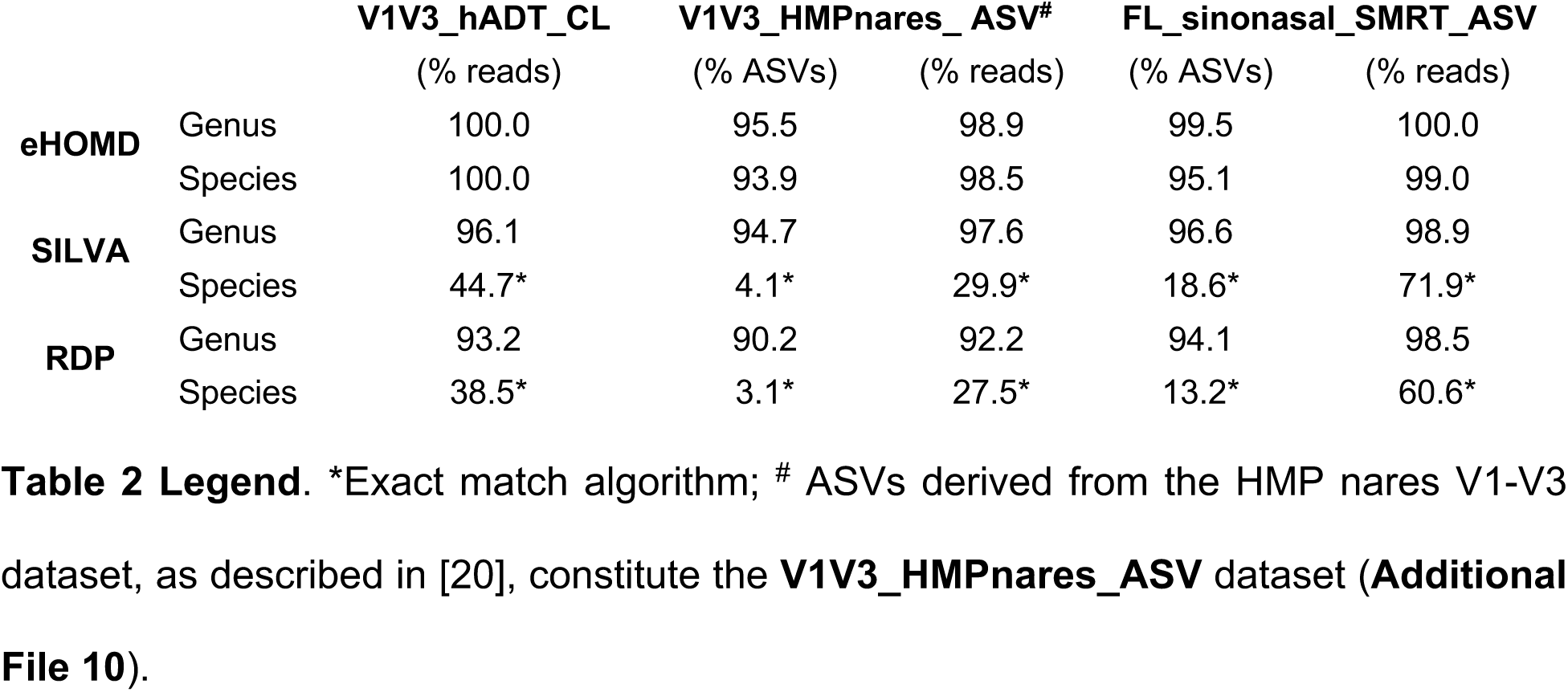
The eHOMD training set is superior for assigning species/supraspecies-level taxonomy to short- and long-read human aerodigestive tract datasets.

A key implication of these data is that our overall method yields comparable species-level results for 16S rRNA gene sequencing of the human aerodigestive tract using V1-V3 short-read sequences, which is very cost effective, compared to using close-to-full-length PacBio SMRT sequences. For example, it readily distinguished sequences of *Staphylococcus aureus* and *Staphylococcus epidermidis*. Another implication is that for species-level analysis of the microbiota of habitats lacking a high-resolution, accurate 16S rRNA gene database, PacBio SMRT sequencing coupled with the newly available DADA2 PacBio pipeline [48] to generate ASVs followed by blastn against NCBI 16S Microbial can provide effective species-level taxonomic assignment (**Table 1**). Of note, the definition of supraspecies is dependent on the database and can vary for different short-read 16S rRNA gene regions. As such, a separate training set needs to be generated for each short-read region of interest. In addition, training sets need regular maintenance in conjunction with the database and need to be regenerated with each major revision of their associated database. In theory, other closed-reference methods for assigning taxonomy could benefit from the addition of an intermediate taxonomic level to preserve the highest level of resolution possible for that method, e.g., suprastrain between species and strain for metagenomics if there are strains that are too closely related to clearly distinguish.

## CONCLUSIONS

Here, we present a systematic approach for generating and validating habitat-specific 16S rRNA gene training sets to achieve species/supraspecies-level taxonomic assignment for short- or long-read 16S rRNA gene sequences. We used the naïve Bayesian RDP classifier with our training set; however, such a training set can be used with other taxonomic classifiers [41, 42, 50-57]. Our training set-construction approach includes several methodological advancements. First, we generated clusters of existing close-to-full length 16S rRNA gene sequences at the 99% level (with ≥ 98% coverage) around highly curated reference sequences. This collection of sequences for each taxon represented the currently known sequence variability for each taxon and increased the accuracy of taxonomic assignment using the naïve Bayesian RDP Classifier (**Figure 3**). Second, we added an intermediate taxonomic level between genus and species, which we dubbed supraspecies, to which we assigned taxa in which sequences overlapped between two or more taxonomic clusters. This increased the percentage of sequences assigned subgenus-level taxonomy by preventing the default of consigning difficult-to-assign sequences to the genus level (**Figure 5**). The process of constructing a short-read training set started with the generation of an effective close-to-full-length training set, which could then be used for taxonomic assignment of long-read 16S rRNA gene sequences such as those from clone libraries and from high-throughput long-read sequencing. As a final critical step, we validated the composition of the eHOMD training set using a PacBio SMRT-sequenced sinonasal dataset consisting of sequences independent of those used to build the training set (**Table 1**). Facilitating species/supraspecies-level analysis of 16S rRNA gene sequence datasets paves the way for cost-effective, population-scale molecular epidemiological microbiota studies that can achieve greater ecological/clinical relevance by reaching species level.

The successful implementation of a training set for species/supraspecies-level assignment of 16S rRNA gene short-read (e.g., V1-V3) datasets involved a collaborative give-and-take to iteratively optimize the sequencing protocol and the analysis workflow in conjunction with each other. As a result, here, we also provided three specific protocol recommendations for sequencing of the 16S rRNA gene V1-V3 region with the Illumina MiSeq. As detailed in **Additional File 4**, first, we show that excellent taxonomic assignment can be achieved with nonoverlapping Illumina reads from V1 and V3 using our recommended sequencing protocol and taxonomic assignment workflow. This counters a common misperception that overlapping Illumina reads are required [6, 58]. The naïve Bayesian RDP Classifier tolerates nonoverlapping V1-V3 sequences; however, some other classifiers might not. Second, we demonstrated the value of initiating Read 1 (R1) from the V3 reverse primer since R1 usually consists of a longer stretch of high-quality sequence. Third, we provide evidence for a role for increased amounts of PhiX when sequencing from the V1 primer, due to the high-degree of sequence conservation 3’ to this primer. This is analogous to the respective recommendations by Illumina and Mitra and colleagues to add PhiX to reach up to or, sometimes, greater than 50% of the total DNA for low diversity samples [59].

We note four limitations to the presented method for constructing a habitat-specific training set. First, capturing the natural sequence variation for the 16S rRNA gene(s) for each taxon is limited by the number of such sequences currently available in public repositories. For example, 140 of the ∼770 taxa in eHOMD are represented by four or fewer distinct close-to-full-length sequences in the training set ***FL_Compilation_TS***, whereas ∼630 taxa are represented by five or greater. Second, the comprehensiveness of a training set depends on the comprehensiveness of the phylogeny-based, habitat-specific database from which it is constructed. Therefore, if an ASV is from a species that is absent from the database, then misclassification of that ASV to the mostly closely related species present in the database is possible. This can also occur during validation and benchmarking if using a simulated dataset with the same origin as the training set [43], e.g., our **V1V3_eHOMDSim_250N100** dataset. This is a known limitation of closed reference-based taxonomic assignment. As such, the known limitations of eHOMD are also limitations of any eHOMD training set [20, 26]. Third, as with databases, training sets also require regular updating over time.

Within these limitations, we note several advantages. First, when coupled with a training set built with our method, the k-mer-based naïve Bayesian approach accommodates the natural variability of 16S rRNA gene sequences that exists within many bacterial species enabling high rates of accurate taxonomic assignment. In contrast, this natural variability limits the utility of any exact match algorithm to assigning species-level taxonomy for only those sequences already existing in a training set (**Table 2**). Second, despite all of the known limitations of a single-gene taxonomic indicator, the huge number of 16S rRNA gene sequences from diverse ecosystems available in public repositories supports the utility of the 16S rRNA gene for taxonomic assignment. In contrast, the utility of WGS metagenomic sequencing, which holds the promise of strain-level taxonomic assignment, remains limited by the quality and comprehensiveness of the genomic database used for closed-reference assignment. For example, at least one single-amplified genome (SAG) of each species is needed for more accurate species-level assignment. This remains problematic for habitats with many as-yet uncultivated species. Also, accurate strain-level assignment is dependent on the presence of SAGs of multiple strains of each species in the reference database. Further, a reference database should be free of chimeric metagenome-assembled genomes (MAGs) that combine genomic sequences that are unique to different strains of a species into one genome.

Finally, there are several additional recommendations to improve taxonomic assignment achieved with our method. First, it is critical to incorporate a stable provisional naming scheme into any habitat-specific database, e.g., the HMT numbers in eHOMD [20, 26]. Second, we recommend validating any training set generated with this method using a habitat-appropriate dataset that is fully independent of the sequences used to construct the training set, as we did with the sinonasal PacBio SMRT-generated dataset. Third, for ASVs that remain unassigned at finer taxonomic levels and are of interest based on their relative abundance and/or prevalence in the population, we recommend two additional steps: first, query these unassigned ASVs against a broad database such as SILVA, RDP and/or NCBI 16S Microbial via blastn (≥98.5% identity over ≥98% coverage) to ascertain if these belong to a named species; and second, if no such named species match exists, then create a new provisional taxon in your database [20, 26]. This approach will identify candidate new taxa, both named and as-yet unnamed, for addition to a habitat-specific database and its associated training set.

## METHODS

### Construction of eHOMD-based training datasets for the naïve Bayesian RDP Classifier

Training datasets were constructed in FASTA format as described on the DADA2 official website (https://benjjneb.github.io/dada2/index.html) [10], with taxonomy in the FASTA header line for each individual sequence. Training sets used in this study are described below.

#### FL_eHOMDrefs_TS

All of the full-length 16S eHOMDrefs, together with their respective taxonomic assignment, were formatted as a training set for the naïve Bayesian RDP Classifier, as above.

#### FL_Compilation_TS

This is the final version of the full-length 16S rRNA gene eHOMD training set, available as **Additional File 2** and for download from eHOMD.org as ***eHOMDv15.1_FL_Compilation_TS.fa.gz***. First, to more precisely calculate the percent identity for recruiting sequences for the training set, we trimmed each of the eHOMDrefs to nucleotides 28 and 1373. This is necessary for two reason: 1) several eHOMDrefs have hanging 5’ or 3’ ends that, if left in place, would affect the calculation of percent identity, and 2) this trimming permitted capture of sequences from the extensive datasets that use the reverse primer at ∼1390. Second, each of the 998 trimmed eHOMDrefs was queried against the NCBI nonredundant nucleotide (nr/nt) database using blastn (NCBI BLAST 2.6.0+ package) (https://www.ncbi.nlm.nih.gov/books/NBK279690/) with the parameters -db nr -remote -perc_identity 97. Nucleotide sequences of GenBank IDs with ≥ 97% sequence identity to any of the eHOMDrefs were downloaded in FASTA format using the efetch.fcgi command of NCBI’s Entrez Programming Utilities (E-utilities; https://www.ncbi.nlm.nih.gov/books/NBK25501/). The blastn hits were downloaded in the same orientation as the eHOMDrefs in two batches: 1) for subject sequence length between 1000 and 2000 nt, the entire subject sequence was downloaded; and 2) for subject sequence length > 2000 bp (e.g., a complete genomic sequence), only the aligned portion of the sequence was downloaded. Sequences < 1000 bp were not downloaded. Third, the 301,794 downloaded sequences, which matched to Human Microbial Taxa (HMTs) at ≥ 97%, were parsed based on their highest sequence percent identity (≥ 99%) and alignment coverage (≥ 98% calculated based on the length of the reference) to any of the eHOMDrefs in a given HMT. The choice of ≥ 99% identity was designed to obtain the centrally conserved set of sequences for each eHOMDref. The choice of ≥ 98% coverage was to ensure that the majority of the close-to-full-length sequence was present. Sequences that matched to multiple HMTs at equal percent identity and coverage were randomly assigned to only one HMT. (The use of supraspecies, see below, mitigates bioinformatic variability introduced at this step.) The ***FL_Compilation_TS*** training set is comprised of a total of 223,144 sequences parsed to their corresponding HMT, with a range of 1 to 4004 sequences per HMT.

#### V1V3_Raw_TS

We generated this training set version in two steps. First, each eHOMDref was individually aligned with the compilation of downloaded sequences that were matched to it at ≥ 99% identity and ≥ 98% coverage (see immediately above). Second, the sequences in the V1-V3 region, defined as positions 40-880 in the gapped eHOMDrefs alignment, were captured and then the alignment gaps removed. These steps were performed using a custom script (**Additional File 11**).

#### V1V3_Curated_TS

Sequences that were identical across V1-V3 in ***V1V3_Raw_TS*** were collapsed into a single sequence with the names of all taxa involved concatenated with a “:” separator. The majority of such concatenations occurred among either the same species, resulting in no name change, or different species of the same genus, resulting in assignment of a concatenated species name. However, there were a number of cases where species from two different genera were involved. These intergenus concatenations were carefully examined case by case. In all but one case, the concatenation was only supported by two to three sequences and, therefore, was deemed unreliable and rejected. After manual examination, only one intergenus concatenation remained. This was between HMT-559 (*Afipia broomeae*) and HMT-597 (*Bradyrhizobium elkanii*) and was supported by 29 sequences. The genus of the concatenated taxa was assigned as *Afipia:Bradyrhizobium* and species as *broomeae:elkanii*. Of note, although these two genera are almost identical on the V1-V3 region, they are 97% identical across the full length of the 16S rRNA gene.

#### V1V3_Supraspecies_TS

This is the final version of the V1-V3 16S rRNA gene eHOMD training set, available as **Additional File 5** and for download from eHOMD.org as eHOMDv15.1_V1V3_Supraspecies_TS.fa.gz (http://www.homd.org/ftp/publication_data/20190709/). The sequences in this training set are the same as those included in the ***V1V3_Curated_TS*** version; only the header information was edited to include the supraspecies notation as a taxonomical level between genus and species. Therefore, instead of the header with seven-levels included in previous versions of the eHOMD training set (i.e., >Kingdom;Phylum;Class;Order;Family;Genus;Species), the ***V1V3_Supraspecies_TS*** training set includes a header with eight levels (i.e., >Kingdom;Phylum;Class;Order;Family;Genus;Supraspecies;Species). A supraspecies was defined between two or more HMTs when there was a phylogenetic distance less than 0.005 between at least one pair of sequences of the corresponding HMTs. To calculate the distances, V1V3 sequences were aligned with MAFTT v6.935b [60] and treed with FastTree version 2.1.9 [61] with default parameters. Pairwise distances between sequences were calculated as the sum of horizontal branch length between two sequence nodes based on FastTree’s default “Jukes-Cantor + CAT” DNA evolution model [61]. The name of the resultant supraspecies is a concatenation of the species names of all HMTs involved, separated by “:” and is assigned at the supraspecies level (seventh level) for all the sequences of the involved HMTs. Sequences that were identical across the V1-V3 region between more than one taxon were assigned the concatenated name both at the supraspecies and species levels (e.g., >Bacteria;Actinobacteria;Actinobacteria;Corynebacteriales;Corynebacteriaceae;Corynebacterium;accolens:macginleyi:tuberculostearicum;accolens:macginleyi:tuberculostearicum). Whereas, the rest of the reads for those taxa were named at the supraspecies level as the concatenated name and maintained their unique identifier at the species level (e.g., >Bacteria;Actinobacteria;Actinobacteria;Corynebacteriales;Corynebacteriaceae;Corynebacterium;accolens:macginleyi:tuberculostearicum;accolens). For taxa not requiring concatenations (see the ***V1V3_Curated_TS*** description), the species designation is repeated at the supraspecies level such that the seventh- and eighth-level designations are identical (e.g., >Bacteria;Firmicutes;Bacilli;Lactobacillales;Carnobacteriaceae;Dolosigranulum;pigrum;pigrum).

### Generation of a V1-V3 16S rRNA gene simulated dataset derived from eHOMD

Simulated reads were generated from each unique sequence in the ***V1V3_Curated_TS*** training set. During read generation, one percent of the bases in each unique initial sequence were randomly selected and changed into a different base to simulate a 1% error rate. The resulting dataset contains 19,480 simulated sequences, each of them labelled with the ID of the parent eHOMDref sequence from which they were derived. Then, each error-simulated read was trimmed into two fragments to simulate Illumina pair-end reads (R1 starting from the V3 primer and R2 starting from the V1 primer). We generated multiple length configurations of both the R1 and R2 fragments. First, fixing R2 at 350 bp, we generated R1 fragments from the V3 primer ranging from 20 to 200 bp (Simulated_R2_350_R1_20-200, **Additional File 12**). Second, fixing R1 at 200 bp, we generated R2 fragments from the V1 primer ranging from 140 to 350 bp (Simulated_R2_140-350_R1_200, **Additional File 13**). Prior to analysis with the naïve Bayesian RDP Classifier, paired reads were connected with a “10N” linker to form one fragment ordered R2-10N-R1 (For further explanation see **Additional File 4**). Subsequent evaluation of the steps for generating the eHOMD training sets in Figures 3 to 5 was done using the configuration R2(250 bp)-10N-R1(100 bp) **(V1V3_eHOMDSim_250N100; Additional File 3**).

### Reanalysis of a sinonasal PacBio-SMRT-sequenced full-length 16S rRNA gene dataset

Earl and colleagues kindly provided the circular consensus sequences (CCS) already demultiplexed, labeled by sample, pooled together and, then, filtered based on length distribution, terminal matches to the primer sequences, not aligning to a provided host or background genome sequence and with a cumulative expected error (EE < 1) [47]. We then used the DADA2 PacBio pipeline to denoise these CCS and identify the relative abundance of each ASV in the dataset; i.e., we used the learnErrors() function with errorEstimationFunction=PacBioErrfun and BAND_SIZE=32 [10, 48]. The sequences of the resulting 204 ASVs are in the **FL_sinonasal_SMRT_ASV** dataset (**Additional File 6**). The V1-V3 region was trimmed using bedtools getfasta with default parameters (bedtools version 2.26.0, https://bedtools.readthedocs.io/en/latest/) and reads were further trimmed into two fragments to simulate Illumina pair-end reads using the configuration R2(250 bp)-10N-R1(100 bp), as described above (**V1V3_sinonasal_SMRT_ASV**; **Additional File 8**).

### Generating a test V1-V3 human aerodigestive tract (hADT) microbiota dataset from full-length 16S rRNA gene clone libraries (CL)

We previously used the close-to-full-length 16S rRNA gene sequences from clone library-based microbiota studies of the human aerodigestive tract, as described in Supplemental Text S1 of [20]: Segre-Kong nostril (SKn) [62-67], Pei-Blaser [68, 69], Harris-Pace [70], van der Gast-Bruce [71], Flanagan-Bristow [72] and Perkins-Angenent [73]. Here, we compiled these into one dataset along with clones from NCBI PopSet UIDs 399192397, 399202217, 399199823, 399197584, 399194446, 399189902, 399186216, 399183739, 399182414, 399179617, 399175646 and 399173254 [74]. Aligned eHOMDrefs (eHOMDv15.1) sequences were trimmed from *E. coli.* position 28-1373 and used to query this compiled dataset via blastn. We retained 27,816 sequences that hit with 100% coverage and ≥ 99.5% identity to 401 HMTs as the full-length <u>h</u>uman <u>a</u>ero<u>d</u>igestive tract clone library dataset (**FL_hADT_CL; Additional File 14**). Of these, 5254 (18.9%) matched to more than one HMT, whereas 22,562 (81.1%) unambiguously matched to single HMTs. Sequences in this full-length CL dataset were then aligned using MAFTT v6.935b with default parameters [60]. Segments corresponding to the V1-V3 region were extracted based on positions of V1-V3 in the alignment (**V1V3_hADT_CL**, **Additional File 9**) using bedtools getfasta with default parameters (bedtools version 2.26.0, https://bedtools.readthedocs.io/en/latest/).

### Taxonomic assignment with the naïve Bayesian RDP Classifier

ASVs and CL sequences were assigned taxonomy using the naïve Bayesian RDP Classifier via the dada2::assignTaxonomy() function in R (an implementation of the algorithm with a k-mer size of 8 and 100 bootstrap iterations) with the specified training set and with outputBootstraps=TRUE and the indicated minBoot value [10, 39]. In addition to our eHOMD training sets, we used the RDP16 (***rdp_train_set_16.fa.gz***) and SILVA132 (***silva_nr_v132_train_set.fa.gz***) genus-level training sets available at https://benjjneb.github.io/dada2/training.html. Since the eHOMD training set ***V1V3_Supraspecies_TS*** generates an output with eight taxonomic levels that might not be compatible with common downstream applications that accept only seven levels, we provide a custom-written R function (**Additional File 15**) that converts the output from eight to seven levels as follows. First, if a sequence is not assigned at species level (i.e., the eighth level) at a given threshold (output as “NA”) then the eighth level is replaced with the classification result and bootstrap value from the corresponding seventh level. Next, the seventh level is deleted and only the resultant merged eighth level that can contain either species or supraspecies information is reported. Please note that this merged last supraspecies-or-species level is labeled simply as species.

### Taxonomic assignment with the DADA2 exact match

ASVs and CL sequences were assigned species-level taxonomy with the RDP16 (***rdp_species_assignment_16.fa.gz***) and SILVA132 (***silva_species_assignment_v132.fa.gz***) training set files downloaded from https://benjjneb.github.io/dada2/training.html using the dada2::assignSpecies() function in R with allowMultiple=TRUE [10].

#### Taxonomic assignment with blastn

The NCBI BLAST 2.6.0+ package (https://www.ncbi.nlm.nih.gov/books/NBK279690/) was installed and the “blastn” command used with max_target_seqs 1, hits with < 98.5% identity were considered nonassigned. ASVs (**FL_sinonasal_SMRT_ASV**) derived from the sinonasal PacBio SMRT-sequenced dataset were also assigned taxonomy using blastn against two databases: the NCBI 16S Microbial database was downloaded from ftp://ftp.ncbi.nlm.nih.gov/blast/db/ on January 2019 [75] and eHOMDv15.1 was converted to a BLAST database using “makeblastdb” from the NCBI BLAST 2.6.0+ package.

## DECLARATIONS

### Ethics approval and consent to participate

All participants provided informed consent and samples used to generate the V1-V3 region 16S rRNA dataset in **Additional File 4 Figure B** were collected under a protocol (#13-14 to FED) approved by the Forsyth Institutional Review Board.

## Consent for publication

Not applicable.

## Availability of data and material

All data are included as additional files. The following are also available for download at eHOMD.org (http://www.homd.org/ftp/publication_data/20190709/): 1) the simulated eHOMD derived dataset (**Additional File 3** as V1V3_eHOMDSim_250N100.fa), 2) both eHOMD training set files (**Additional File 2** as eHOMDv15.1_FL_Compilation_TS.fa.gz; and **Additional File 5** as eHOMTv15.1_V1V3_Supraspecies_TS.fa.gz) and 3) both custom scripts (**Additional File 11** as retrieveSegment.py and **Additional File 15** as eight2seven.R).

## Authors’ contributions

Conceived Project: YH, IFE, FED, KPL. Designed Project: YH, IFE, FED, KPL. Generated and Analyzed data: YH, IFE, TC, ML, AK. Interpreted results: YH, IFE, TC, ML, AK, FED, KPL. Generated figures and tables: IFE, YH. Wrote manuscript: KPL, IFE. All authors reviewed and edited the manuscript and approved the final version.

## Acknowledgements

We are grateful to Joshua Earl, Joshua Chang Mell and colleagues for providing the intermediate data table that facilitated analysis with a newly available algorithm and to members of the Lemon Lab and the Starr-Dewhirst-Johnston-Lemon Joint Group Meeting for helpful questions and suggestions throughout the project.

## ADDITIONAL FILES.

**Additional File 1**: The expected effect of an uneven distribution of sequences among taxa in a training set for the Naïve Bayesian RDP Classifier.

**Additional File 2**: eHOMDv15.1_FL_Compilation_TS.fa. The 16S rRNA gene full-length eHOMD training set (***FL_Compilation_TS***).

**Additional File 3**: V1V3_eHOMDSim_250N100.fa. The simulated eHOMD-derived **V1V3_eHOMDSim_250N100** dataset.

**Additional File 4**: A method for achieving highly informative 16S rRNA gene V1-V3 region sequencing data using Illumina MiSeq.

**Additional File 5**: eHOMDv15.1_V1V3_Supraspecies_TS.fa. The 16S rRNA gene V1-V3 eHOMD training set (***V1V3_Supraspecies_TS***).

**Additional File 6**: FL_sinonasal_SMRT_ASV.fa. Sequences of the resulting 204 ASVs in the Earl-Mell sinonasal 16S rRNA gene dataset (**FL_sinonasal_SMRT_ASV** dataset).

**Additional File 7**: Earl-Mell_sinonasal_analysis.xlsx with legend and tabs A-D. Species-level taxonomic assignment of the Earl-Mell sinonasal 16S rRNA gene dataset.

**Additional File 8**: V1V3_sinonasal_SMRT_ASV.fa. Sequences of the resulting 204 ASVs in the **FL_sinonasal_SMRT_ASV** dataset trimmed to the V1-V3 region (**V1V3_sinonasal_SMRT_ASV** dataset).

**Additional File 9**: V1V3_hADT_CL.fa. V1-V3 trimmed 16S rRNA gene <u>h</u>uman <u>a</u>ero<u>d</u>igestive tract clone library dataset (**V1V3_hADT_CL** dataset).

**Additional File 10**: V1V3_HMPnares_ASV.fa. ASVs derived from the HMP nares 16S rRNA gene V1-V3 dataset (**V1V3_HMPnares_ASV** dataset).

**Additional File 11**: retrieveSegment.py. Custom script developed to generate a trimmed version of a training set (in our example to positions 40-880 in the gapped eHOMDrefs alignment) from a compilation of 16S rRNA gene full-length sequences.

**Additional File 12**: Simulated_R2_350_R1_20-200.7z. Simulated eHOMD-derived dataset versions with R2 fixed at 350 bp and R1 fragments from the V3 primer ranging from 20 to 200 bp.

**Additional File 13**: Simulated_R2_140-350_R1_200.7z. Simulated eHOMD-derived dataset versions with R1 fixed at 200 bp and R2 fragments from the V1 primer ranging from 140 to 350 bp.

**Additional File 14**: FL_hADT_CL.fa. Full-length 16S rRNA gene <u>h</u>uman <u>a</u>ero<u>d</u>igestive tract clone library dataset (**FL_hADT_CL** dataset).

**Additional File 15**: eight2seven.R. Custom-written R function that converts the dada2::assignTaxonomy() output from eight to seven taxonomic levels for compatibility with downstream applications.

