## Supplemental File 1 for "Construction of habitat-specific training sets to achieve species-level assignment in 16S rRNA gene datasets"

**Additional File 1: The expected effect of an uneven distribution of sequences among taxa in a training set for the Naïve Bayesian RDP Classifier**

As illustrated in **Figure 2,** the naïve Bayesian RDP Classifier algorithm [1] works as follows. Given a training dataset that contains N unique sequences and in which the word wi appears n(wi) times, a prior is calculated so Pi = (n(wi)+0.5)/(N+1). Then, for a given taxon (T) that contains M sequences and in which the word wi appears m(wi) times, the taxon-specific conditional probability for wi is P(wi|T) = (m(wi) + Pi)/(M + 1). For a sequence (S) containing many words, w1,w2,w3, …,wj, the joint probability that S is derived from T is P(S|T) = P(w1|T)*P(w2|T)*P(w3|T)*…* P(wj|T). As M increases, the estimation of the prior improves, and the resolution (i.e., granularity) improves because now P(wi|T) = (m(wi) + Pi)/(M + 1) is no longer limited to only two possible values, (1+Pi)/2 and (0+Pi)/2. More granular changes in P(wi|T) allow sequences with smaller differences to be distinguished, thus resolution improves.

When there is a disproportionally high number of sequences from a particular taxon present in the training set, would sequences get disproportionately assigned to that taxon? We observed that more sequences were classified unambiguously as the number of sequences (M) for each taxon in the training set increased (**Figure 3**). This is consistent with what Werner and colleagues reported [2]. To explain this, we will use a simplified example with two training sets: Set1 contains two taxa, A and B, with only one reference sequence each; while Set2 contains 90 sequences for taxon A and 10 sequences for taxon B, i.e., there is an uneven distribution of reads across taxa in Set2. In this simple example both the sequence of taxon A and taxon B contain four words. Three of the words, w1, w2 and w3 are shared but the fourth one is different, being w4 for taxon A and w5 for taxon B, as shown below:

Taxon A reference: w1w2w3w4

Taxon B reference: w1w2w3w5

Here we can see that for words shared by different taxa, the number of sequences in the training set and how well each taxon is represented have minimal effect. If we analyze the probabilities for the common words (w1, w2 and w3) the two different training sets yield similar results, i.e., in both cases P(wi|A) = P(wi|B):

Set1: taxon A (1 sequence) + taxon B (1 sequence)

Pi = (2 + 0.5) / (2+1) = 0.83

P(wi|A) = P(wi|B) = (1+0.83) / (1+1) = 0.915

Set2: taxon A (90 sequences) + taxon B (10 sequences)

Pi = (100 + 0.5) / (100+1) ~= 1

P(wi|A) = (90 + Pi) / (90+1) ~= 1

P(wi|B)= (10+Pi) / (10+1) ~= 1

In contrast, words that distinguish taxon A from taxon B (w4 and w5) have a large effect, where w4 is only in A and w5 is only in B. As shown below, using Set2 increases the difference between P(wi|A) and P(wi|B) and, thus, improves the ability to classify to either A or B.

Set1: taxon A (1 sequence) + taxon B (1 sequence)

Priors P4 = P5 = (1+0.5) / (1 + 1) = 0.75

P(w4|A) = (1+0.75) / (1+1) = 0.875

P(w4|B) = (0+0.75) / (1+1) = 0.375

P(w5|A) = (0+0.75) / (1+1) = 0.375

P(w5|B) = (1+0.75) / (1+1) = 0.875

Set2: taxon A (90 sequences) + taxon B (10 sequences)

Prior P4 = (90+0.5) / (100 +1) = 0.896

P(w4|A) = (90+0.896) / (90+1) = 0.999

P(w4|B) = (0+0.896) / (10+1) = 0.081

Prior P5 = (10+0.5)/(100+1) = 0.104

P(w5|A) = (0+0.104)/(90+1) = 0.001

P(w5|B) = (10+0.104)/(10+1) = 0.919

To be more specific, for this example, when there are two query sequences, SA and SB, their probability of belonging to taxon A or taxon B is calculated as the following:

P(SA|A) = P(w1|A)*P(w2|A)*P(w3|A)*P(w4|A)

P(SA|B) = P(w1|B)*P(w2|B)*P(w3|B)*(Pw4|B)

P(SB|A) = P(w1|A)*P(w2|A)*P(w3|A)*P(w5|A)

P(SB|B) = P(w1|B)*P(w2|B)*P(w3|B)*(Pw5|B)

We should be able to classify SA as being derived from taxon A because P(SA|A) > P(SA|B) and SB to be derived from taxon B because P(SB|A) < P(SB|B), and it is obvious that Set2, with 90 sequences for taxon A and 10 sequences for taxon B, provides better distinction.

For simplicity, the example discussed above assumed that all 90 sequences of taxon A contain w4 but not w5 and all 10 sequences of taxon B contain w5 but not w4. This was only to illustrate that imbalance between the number of sequences included from each taxon does not bias the classification. This is precisely because P(wi|T) = (m(wi) + Pi)/(M+1). When m(wi) increases, so does M, thus P(wi|T) should be relatively stable regardless of number of sequences collected. The only concern would be that there might be situations where the proportion of sequences in M containing wi does not reflect the real proportion in nature. However, we think any proportion is likely to give a better estimate than 1 or 0, which is the case when you only have one sequence for each taxon.
