## Supplemental File 4 for "Construction of habitat-specific training sets to achieve species-level assignment in 16S rRNA gene datasets"

**Additional File 4: A method for achieving highly informative 16S rRNA gene V1-V3 region sequencing data using Illumina MiSeq.**

The length of the V1-V3 16S rRNA gene region, which corresponds to *E. coli* positions 28 to 517, is highly variable in bacteria. The eHOMD includes 770-plus species-level taxa and the eHOMD phylogeny is available at http://www.homd.org/ftp/HOMD_phylogeny/current/. For all but three taxa in eHOMD, the V1-V3 region size ranged from 431 to 523 base pairs (bp). Three Campylobacter species, with intervening sequences in the V2 region, were up to 653 base pairs in length (**Figure A.i, Additional File 4**). This size variability makes it extraordinarily difficult to capture full-length V1-V3 sequences for all bacterial taxa using Illumina technology. Even using the 500 or 600 cycle MiSeq kits, this is impeded by the decline in sequence quality as the run progresses (**Figure A.i, Additional File 4**). Many analysis pipelines reject read pairs that do not overlap. Excluding taxa with longer V1-V3 regions biases results and generates a false understanding of these microbial communities. To solve these problems, we hypothesized that there are specific subregions within V1-V3 that are most informative such that complete sequencing across the V1-V3 region via overlapping paired reads is unnecessary. To identify the more informative V1-V3 subregions, we evaluated Shannon entropy for each position across all of taxa represented by eHOMD reference sequences (eHOMDrefs) (**Figure A.ii, Additional File 4**). Based on examination of the entropy values across the V1-V3 region, we postulated that sequencing 300 bp from the V1 primer and 150 bp from the V3 primer would capture the majority of the sequencing diversity needed to maximize species-level taxonomic assignment to the V1-V3 region for species included in eHOMD.

Moreover, one advantage of the naïve Bayesian RDP Classifier, which is a k-mer-based Bayesian approach, is that it can provide a single taxonomic assignment when analyzing

Illumina pair-end data without requiring overlap of the forward and reverse reads. Therefore, from each primer, V1 and V3, we separately visualized at what read length there was no longer a gain in the percentage of classified sequences using a simulated eHOMD-derived dataset. To accomplish this for primer V3, we fixed the read length from primer V1 at 350 bp (**Figure A.iii, Additional File 4**) based on the indication in Figure Aii that the first 300 bp captures the majority of sequence variability with the extra 50 bp allowing for variability in region length. Similarly, for primer V1, we fixed the read length from primer V3 at 200 bp (**Figure A.iv, Additional File 4**). We then determined the percentage of sequences classified at species level at increasing read lengths from the complementary primer using the 998 eHOMDrefs as a training set (***FL_eHOMDrefs_TS***). Based on these assays, species-level assignment plateaued across all bootstrap values at 70 bp from primer V3 (**Figure A.iii, Additional File 4**). Moreover, species-level assignment started to plateau across all bootstrap values around 210 bp from primer V1 (**Figure A.iv, Additional File 4**). These results indicated that species-level identification can be achieved with nonoverlapping reads for the 16S rRNA gene V1-V3 region using the naïve Bayesian RDP Classifier, at least for the taxa in eHOMD.

Furthermore, these results establish guidance for actual read lengths needed for Illumina sequencing of the V1-V3 16S rRNA gene region. Knowing this, we next determined experimentally what length of high-quality sequences are obtainable using primers V1 and V3 with 500 cycles on the Illumina MiSeq. Amplicon-based libraries can be challenging to sequence using Illumina's 2-channel sequencing chemistry since their low diversity hinders correct identification of DNA clusters on the Illumina-flow cell and accurate base calling [1]. To mitigate this, we changed Read 1 (R1) to start from the V3 primer instead of V1, since clusters are defined very early in an Illumina run (first 4 cycles) [2] and sequences are mostly identical in the first positions immediately 3' of the V1 primer. We also increased the proportion of PhiX from a range of 10-25% up to a range of 30-50%, as recommended in [2]. Together, these steps accomplished a significant improvement in sequence quality (**Figure Bi-iii, Additional File 4**).

During our experiments, we also observed that variability in library quality, e.g., from low biomass habitats such as the human nostrils, can negatively impact the accuracy of library quantification leading to an inadvertently low percentage of PhiX. Because the informative region from V1 (R2) includes the first 250 bp from the primer (**Figure A.iv, Additional File 4**), we routinely perform asymmetrical sequencing of R1 and R2 with R1 = 100 bp and R2 = 400 bp (e.g., **Figure B.iii, Additional File 4**) to avoid having the decrease in sequence quality that is sometimes observed at the end of an Illumina MiSeq run falling within the informative region from the V1 primer (R2) (**Figure B.ii, Additional File 4**). After trimming poor quality regions from each experimentally-derived read, this generally results in high-quality sequences of 200 to 250 bp from primer V1 and 100 bp from primer V3. Thus, for the simulated eHOMD-derived dataset (**V1V3_eHOMDSim_250N100, Additional File 3**) created to test each step in the development of the eHOMD training set, we mimicked these Illumina-generated nonoverlapping reads by designing artificial reads consisting of 250 bp from primer V1 and 100 bp from primer V3. To enhance clarity, the names of datasets appear in bold whereas the names of training sets appear in bold italics.

**Methods for Additional File 4**

**Determination of the 16S rRNA gene V1-V3 sequence length for each taxon in eHOMD.** To construct Figure A.i, the 998 eHOMDv15.1 aligned 16S reference sequences (eHOMDrefs) were downloaded in fasta format from eHOMD (http://www.homd.org/?name=seqDownload&type=R) [3]. For each of the ~770 Human Microbial Taxa (HMTs) in eHOMD, one eHOMDref was selected as a representative of that given taxon. The resulting alignment was trimmed from *E. coli* position 28 to 517 (V1-V3 region) and further trimmed to positions 28-220 (V1-V2 region) and 436-517 (V3 region).

**Entropy calculation for the 16S rRNA gene V1-V3 region for each taxon in eHOMD.** To construct Figure A.ii, sequences in the eHOMDv15.1 alignment were trimmed from aligned positions 40 to 880. The resulting sequences with alignment gaps are much longer than actual V1-V3 sequences. Shannon entropy for each position in the aligned sequences was estimated with a maximum likelihood method using the "entropy" R package [4]. Because the distribution of ungapped V1-V3 lengths centered around 489 bases, a V1-V3 fragment with an ungapped length of 489 (eHOMDref 762_5043) was randomly chosen from the V1-V3 sequence collection as a representative onto which the entropy results were plotted.

**Nonoverlapping V1-V3 16S rRNA gene sequencing.** Supragingival pooled plaque samples were deposited into collection tubes containing 150 µL TE buffer. Samples were incubated at 37°C for 3 hrs with 1 µL of Lucigen's Ready-Lyse Lysozyme, and then subsequently processed using Lucigen's MasterPure Complete protocol beginning at step C-Lysis of Fluid or Tissue Samples. The method deviated from the protocol at step 4 where the incubation time was increased to 30 mins and steps 5 and 6 were omitted. Samples were precipitated using step F-Precipitation of Total Nucleic Acids and were resuspended in 25 µl of TE buffer instead of the suggested 30 µl. 16S rRNA gene libraries were prepared by targeted amplicon sequencing performed using a custom dual-index protocol [5]. The 16S rRNA gene primers described by Allen and colleagues (**Table A, Additional File 4)** were used to amplify the V1-V3 region of the 16S rRNA gene [6]. These primers provide broad coverage of 16S rRNA genes while maintaining high sensitivity. The sample libraries were prepared using a 22-cycle PCR reaction to reduce the chimera formation that is common when using a high cycle number. The final

PCR products were purified using Ampure XP beads, pooled in equal amounts, and gel purified using the QIAGEN MinElute Gel Extraction Kit. Purified, pooled libraries were quantified using the NEBNext Library Quant Kit for Illumina. Final libraries were sequenced on Illumina® MiSeq™ with a v2 reagent kit (500 cycles) using a 250x250 or 100x400 configuration. The sequencing was performed at a 10 pM loading concentration with the indicated percentage of PhiX added. Quality Phred Score of the reads was assessed using FastQC [7].

**Tables for Additional File 4**

**Table A. Sequencing primers used to amplify the V1-V3 region of the 16S rRNA gene.**

| |
|---|
| **V1V3 Region PCR Primers (Allen et al., 2016)**<br><br>518F (forward primer):<br>AATGATACGGCGACCACCGAGATCTACAC ATCGTACG TATGGTAATT**CAATTACCGCGGCTGCTGG**<br>27R (reverse primer):<br>CAAGCAGAAGACGGCATACGAGAT AACTCTCG AGTCAGTCAG**CCGAGTTTGATCMTGGCTCAG**<br><br>Legend:<br>Illumina 5' or 3' adaptor<br>Barcode<br>Primer Pad Region (linker)<br>**16S Primer** |
| **Sequencing Primers**<br><br>Read 1 Sequencing Primer (V3_518):<br>TATGGTAATTCAATTACCGCGGCTGCTGG<br>Read 2 Sequencing Primer (V1_27):<br>AGTCAGTCAGCCGAGTTTGATCMTGGCTCAG<br>Index Sequencing Primer:<br>CTGAGCCAKGATCAAACTCGGCTGACTGACT |

**Figures for Additional File 4**

**Figure A. 16S rRNA gene V1-V3 region sequences do not require overlap to provide maximal information for human aerodigestive tract-associated bacteria.** (**i**) Rank order of eHOMD taxa based on the nucleotide length of regions V1-V3 (purple), V1-V2 (orange) and V3 of the 16S rRNA gene. (**ii**) Shannon Entropy (H) across the 16S rRNA gene V1-V3 region for all taxa in eHOMD. For easier visualization, bars are color-coded in gray scale based on their entropy values, i.e., the taller a bar is the darker it is. The percentage of eHOMD-derived simulated reads that can be classified with the ***FL_eHOMDrefs*** training set at bootstrap values from 70 to 100 (see key) using (**iii**) a fixed read length of 350 bp from primer V1 and variable read lengths from primer V3 or (**iv**) a fixed read length of 200 bp from primer V3 and variable read lengths from primer V1.

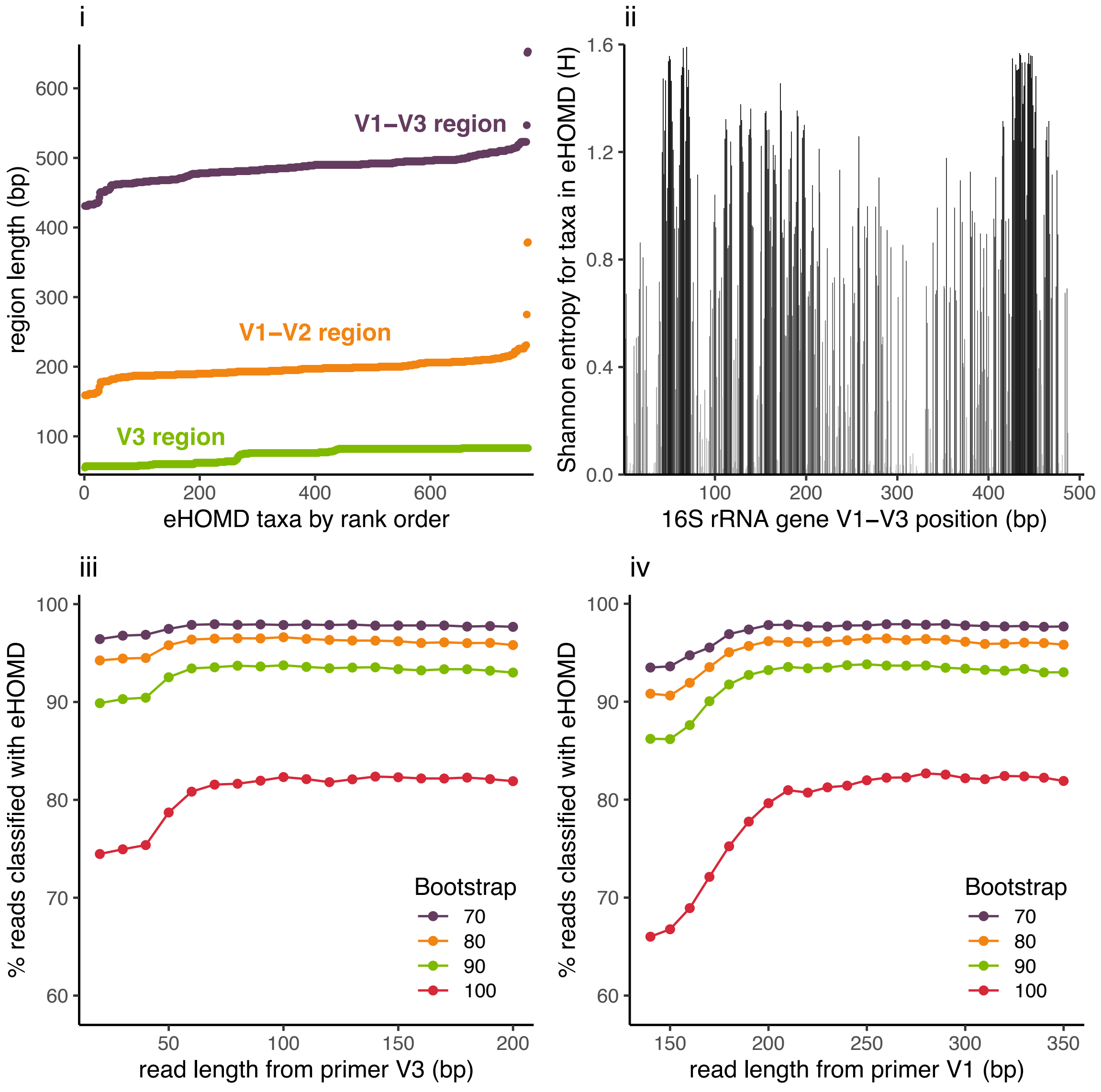

**Figure B. Quality Phred score plots of representative 16S rRNA gene V1-V3 region Illumina runs.** (**i**) Symmetrical sequencing run (R1 = 250 nt and R2 = 250 nt) and PhiX= 20%. (**ii**) Symmetrical sequencing run (R1 = 250 nt and R2 = 250 nt) and PhiX= 34%. (**iii**) Asymmetrical sequencing run (R1 = 100 nt and R2 = 400 nt) and PhiX= 47%.

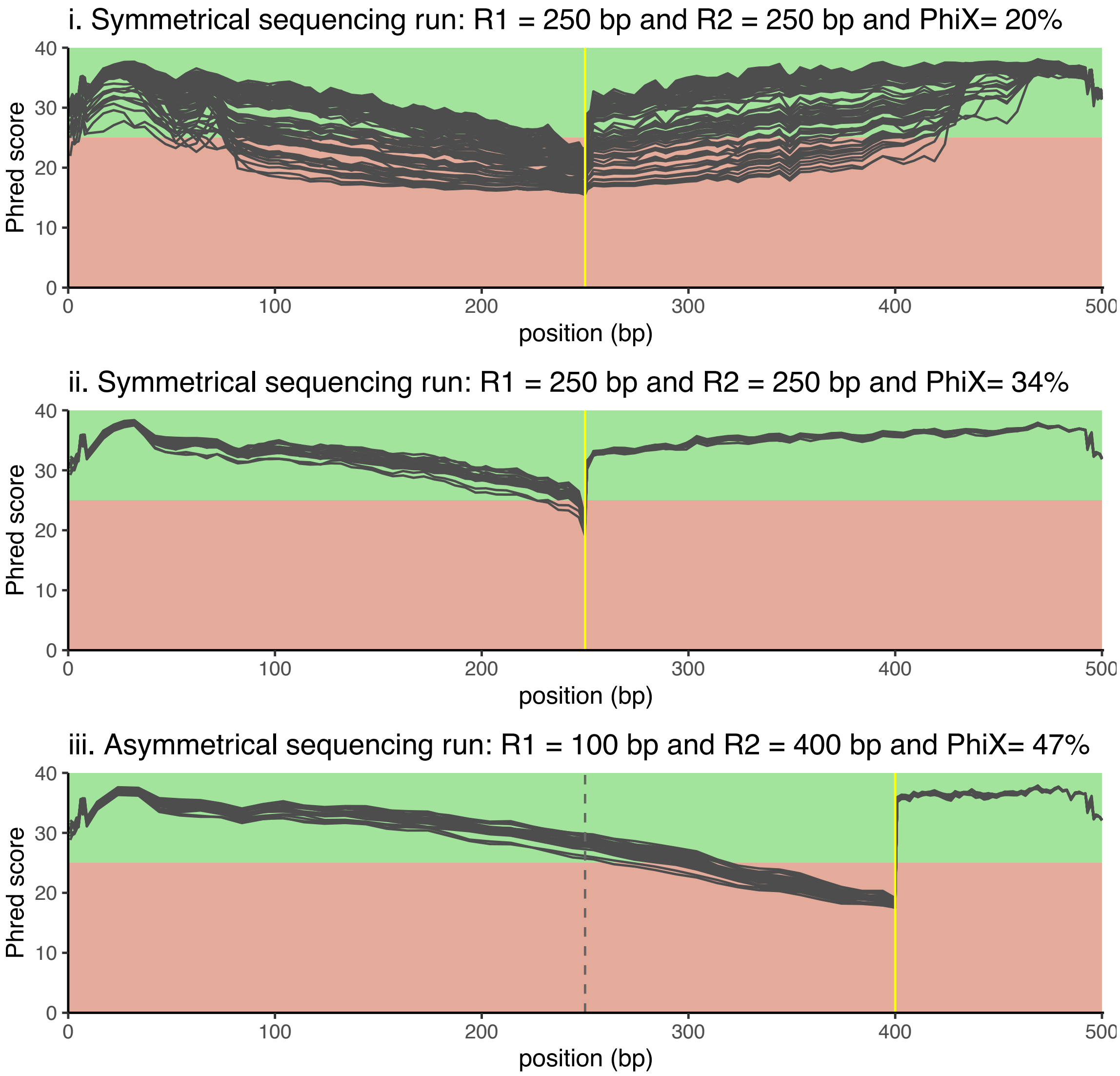
